# Optimizing Digital PCR Assays for Multiplexed Microbial Source Tracking in Surface Waters

**DOI:** 10.1101/2024.09.07.611803

**Authors:** Shimul Ghosh, Aaron Bivins

## Abstract

Digital PCR (dPCR) shows great promise for precise, sensitive, and inhibition-resilient measurement of nucleic acids in surface water, providing an advantage for applications such as microbial source tracking (MST). Herein, we describe our empirical optimization of two triplex format dPCR reactions (Triplex 1 - Cow M2, Rum2Bac, Cow M3; Triplex 2 - HF183/BacR287, Pig2Bac, Entero1a) on the QIAcuity One system for MST. Each triplex produced gene copy measurements similar to single-plex format assays for a standard reference material (SRM 2917) at a fixed concentration and along a concentration gradient. For achieved water samples previously tested by single-plex assays, each triplex also produced MST marker measurements comparable to the single-plex results. The triplexes described here can be directly adopted for MST on the QIAcuity, or the optimization protocol we demonstrate can be used to develop additional multiplex assays on the QIAcuity system.

## 1 Introduction

Digital PCR (dPCR) makes use of PCR reaction partitioning and the Poisson distribution to quantify nucleic acid templates using a most probable number (MPN) approach. Unlike quantitative PCR (qPCR), dPCR does not require the preparation and analysis of calibration curves and standards to produce a quantitative result. Instead, dPCR estimates the number of template copies by measuring the presence or absence of the target in a large number of independent PCR partitions and then estimating the copy number via a Poisson model (***1***). Although digital PCR was conceptualized before qPRC, the nano-scale partitioning required for precise quantification was not operationalized in a reliable format until 1998 (***2, 3***). The first commercially available dPCR system was brought to market by Fluidigm in 2006 (***4***). In 2011, BioRad Laboratories introduced the QX100, which revolutionized dPCR via a reliable water-oil emulsion process that made the dPCR format more accessible to research laboratories (***5***).

Digital PCR saw immediate applications in clinical cell biology, particularly for the sensitive detection and quantification of rare mutations and pathogens in the fields of oncology and pathology. However, the sensitive and calibration-curve-free dPCR format also offers improved resilience to inhibitors, which is advantageous for environmental microbiology, particularly water microbiology, where molecular measures are becoming increasingly common (***6, 7***). Recently, the uptake of dPCR for water microbiology has been accelerated by the widespread use of wastewater testing for public health surveillance during the COVID-19 pandemic, where reliability is crucial to inform decision-making (***8, 9***). Another compelling use case for dPCR is microbial source tracking (MST). During MST, molecular assays are designed to be specific to particular sources of fecal contamination and used to test environmental water samples to inform risk management in recreational waters (***10***). The U.S. EPA has developed qPCR-based methods to measure human-specific fecal contamination (HF183/BacR287) and fecal indicator bacteria Enterococci (Entero1a) (***11, 12***).

We adapted these two TaqMan probe assays from the U.S. EPA, along with four additional MST assays, to dPCR format and optimized two triplex reactions for simultaneous measurement of fecal contamination from ruminants, pigs, cows, and humans in surface waters (Table 1). Each optimized triplex provided precise quantification of a standard reference material that was statistically similar to singe-plex measurements for both a fixed concentration (Figure 4) and a concentration gradient (Figure 5). Each triplex also produced qualitative and quantitative MST results that were comparable to previous single-plex testing of DNA extracts from surface water samples (Figures 8 and 9). Because dPCR multiplexing development is not thoroughly described in the peer-reviewed literature, here we describe our multistep protocol to optimize and validate triplex reactions in dPCR format. Directly, the triplexes we have optimized can be replicated for MST in surface waters using the QIAcuity dPCR system. Indirectly, the approach we have described could be used to optimize and validate other multiplex assays in dPCR format for applications in water microbiology.

**Table 1.**
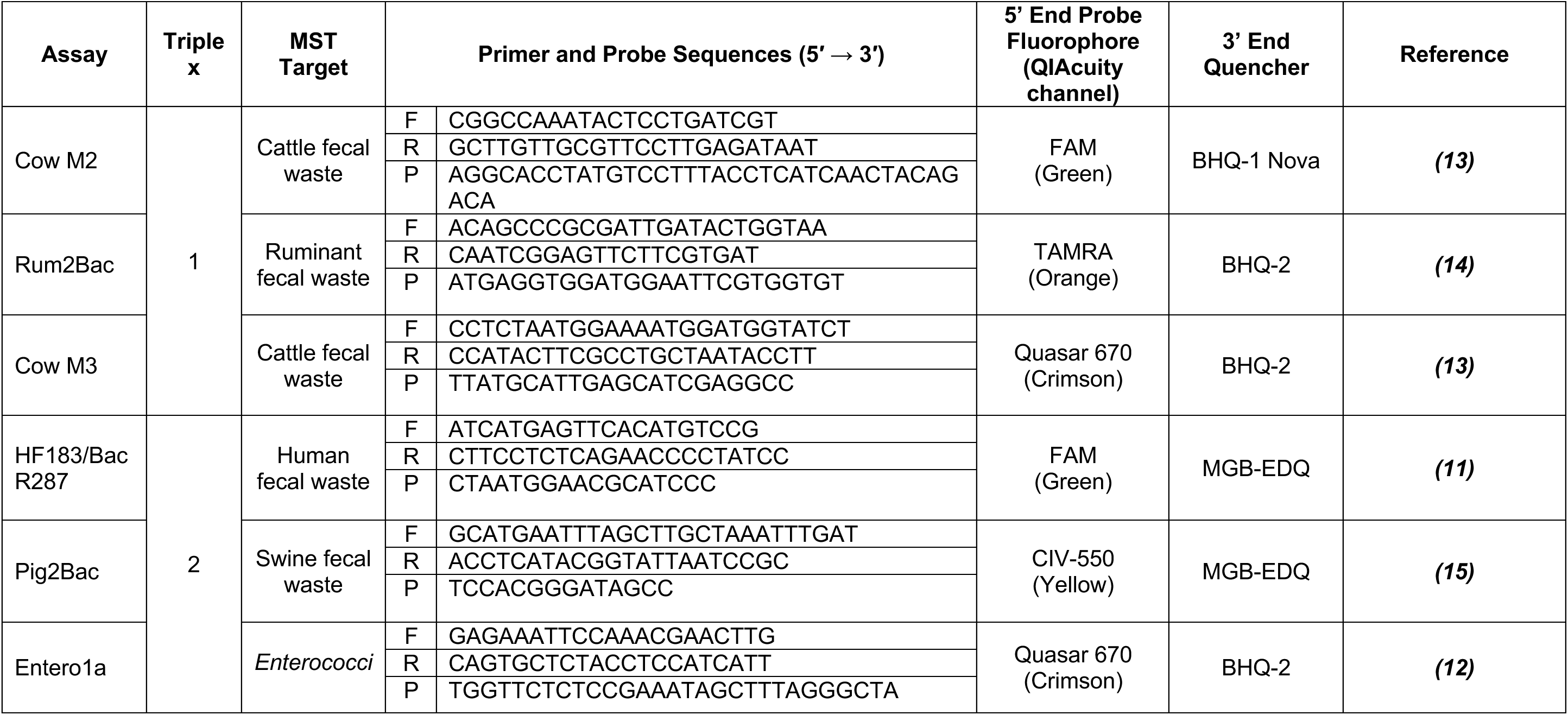
Primer sequences, probe sequences, and fluorophore and quenchers for the MST assays as adapted to dPCR.

## 2 Materials

Prepare all dPCR reaction solutions, primer, probe, and control material dilutions with nuclease-free molecular-grade water and/or TE (Tris-EDTA) buffer at pH 8.0. To maximize precision and comparability between dPCR experiments, use calibrated pipettes with proper technique for all liquid handling described herein. To minimize the opportunity for cross-contamination, all dPCR pipetting should be performed in a nucleic acid workstation (dead air box) using barrier tip pipettes with aseptic techniques appropriate for the relevant biosafety level of the laboratory (BSL 2 in our case). All surface water samples should be handled in a biosafety cabinet per the approved biosafety protocols to minimize the probability of incidental infection among laboratory personnel. Diligently follow all waste disposal regulations and procedures when disposing of waste materials.

### 2.1 Diluents, Specialty Plastic Ware, and General Lab Equipment

1. TE (Tris-EDTA) Buffer, pH 8.0 (VWR, E112-100ML)
2. Molecular-grade nuclease-free water (VWR, VWRL0201-0500)
3. 1.5 mL DNA LoBind tubes (Eppendorf, 022431021)
4. Snap-Cap 5mL centrifuge tubes (VWR, 10002-731)
5. 1.7 mL microcentrifuge tubes (VWR, 87003-294)
6. 0.2 mL PCR strip tubes (Axygen, PCR-0208-AF-C)
7. Microcentrifuge (VWR, 76269-064)
8. Refrigerated benchtop centrifuge 5810R (Eppendorf, 022625501)
9. Centrifuge rotor A-4-62 (Eppendorf, 022638009)
10. Analog Vortex Mixer (VWR, 10153-838)

### 2.2 Primers and probes for dPCR

Digital PCR assays for each MST marker were adapted from published qPCR TaqMan probe assays (Table 1). The primers for each assay were purchased as custom single-stranded DNA oligonucleotides from Integrated DNA Technologies. Primers were ordered as “LabReady” (100 µM stock solution suspended in IDTE buffer at pH 8.0). From the “LabReady” stock solutions, 10 µM working solutions were prepared for each primer in a 1.7 mL microcentrifuge tube using an aliquot of the stock and molecular-grade water.

Multiplexing dPCR assays requires careful consideration of probe fluorophore selection to minimize cross talk between dye channels. The probes for each the dPCR assays were purchased as custom DNA probes from Biosearch Technologies (Note 1). For assays requiring a minor groove binder (MGB) (HF183/BacR287, Pig2Bac), we used Biosearch’s MGB probes, which feature an MGB moiety conjugated to an Eclipse Dark Quencher (EDQ). For the Cow M2 assay, which has a probe length greater than 25 bases, we used Biosearch’s BHQnova™ probe that uses both a Black Hole Quencher (BHQ™) and an internal Nova quencher to increase quenching efficiency. For the remaining three assays (Entero1a, Cow M3, Rum2Bac), we used Biosearch’s BHQ™ probes with fluorophores and quenchers selected per their recommendations (Note 2). We selected the fluorophores for each probe based on the anticipated assay groups for Triplex 1 and Triplex 2, as summarized in Table 1. Upon receipt, all probes were resuspended to 100 µM in TE (Tris-EDTA) buffer at pH 8.0 per the manufacturer’s recommendation. We then created a 25 µM working solution for each probe in a 1.7 mL microcentrifuge tube using an aliquot of the 100 µM stock solution and molecular-grade water. All stock and working solutions for primers and probes were stored in a reagent-specific refrigerator at -20 C until use.

### 2.2 Control plasmid

Standard Reference Material 2917 Plasmid DNA for Fecal Indicator Detection and Identification (National Institute for Standards and Technology, SRM 2917) (Note 3). SRM 2917 was stored at 4 C when not in use per NIST’s specification.

### 2.3 Purified Nucleic Acids from Surface Water Samples

To test the performance of the triplex reactions on environmental material, we made use of six archived nucleic acid extracts derived from surface water samples collected and tested during previous MST projects. Briefly, these extracts were produced by filtering 50 to 100 mL of surface water through a polycarbonate membrane (Millipore, HTTP04700). The membrane was then aseptically rolled and placed in a PowerWater DNA Bead Tube (Qiagen, 14900-50-NF-BT). DNA was then extracted from the filter using the DNeasy PowerWater Kit (Qiagen, 4900-100-NF) following the manufacturer’s protocol. Following the initial testing for a set of MST markers, the purified DNA was stored at -80 C until interrogation via dPCR using the triplex assays described herein.

### 2.4 dPCR system and reagents

1. QIAcuity One, 5plex dPCR System (Qiagen, 911022)
2. QIAcuity Probe 5 mL PCR Kit (Qiagen, 250102)
3. QIAcuity Nanoplate 26k 24-well (Qiagen, 250001)
4. QIAcuity Nanoplate 26k 8-well (Qiagen, 250031)
5. Nanoplate Tray (Qiagen, 250098)
6. Nanoplate Seals (Qiagen, 250099)
7. QIAcuity Suite Software v. 2.5.0.1
8. QIAcuity Control Software v. 2.5.0.24
9. QIAcuity Volume Precision Factor v. 5.0 (VPFV50SOW891005)

## 3 Methods

All dPCR experiments, including software setup, reaction setup, plate sealing, plate loading, partitioning, thermal cycling, and imaging, were performed following the general protocols and procedures detailed in the QIAcuity User Manual, QIAcuity Application Guide, and QIAcuity Probe PCR Kit (***16–18***). Prior to beginning the reaction setup and plate loading, as described in the sections below, the QIAcuity One dPCR system should be started, and the QIAcuity Control and Suite software should be initiated to ensure the instrument is in good working order. It is also best to configure the experimental run in the Suite Software prior to dispensing reagents. This will ensure any hardware and/or software issues are identified and resolved prior to expending reagents or consumables. For the sake of brevity, we have not detailed every standard operating procedure for the QIAcuity system. However, we have noted and detailed any deviation from the standard procedures in the sections to follow.

### 3.1 Single-plex Assay Primer/Probe and Annealing Temperature Optimization

We began our optimization protocol by examining the performance of each MST assay in single-plex format for combinations of annealing temperature (55, 57, 60 C) and primer/probe concentration (S1, S2, S3). These parameters have been previously identified as important for optimizing dPCR performance (***19***). The goal of these single-plex experiments is to ensure reliable separation between the positive and negative dPCR partitions at an annealing temperature that is compatible with the other assays in the proposed triplex and examine the consistency of the separation with different primer/probe concentrations (Note 4).

1. **SRM 2917 working solution preparation.** Based on the anticipated sample size for the single-plex and duplex experiments, a 2,500 µL SRM 2917 working solution at a final concentration of approximately 10,000 GC/µL was prepared by pipetting 2,453.8 µL of molecular-grade water into a Snap-Cap 5mL centrifuge tube followed by 46.2 µL of SRM 2917 Level 6 stock solution (certified by NIST at 541,286 +/-7,700 GC/µL). The SRM 2917 working solution was stored at 4 C when not in use for the duration of the single-plex and duplex experiments (9 experimental days performed over 22 calendar days). Prior to any subsampling for dPCR experiments, the tube was vortexed for 30 seconds and briefly spun down in a centrifuge.
2. **dPCR super mix preparation.** Single-plex dPCR optimization experiments were performed for one assay each day by running three different dPCR super mixes (S1, S2, S3) on a single QIAcuity Nanoplate 26k 24-well plate (one reaction mixture per 8-well column) with a single annealing temperature during thermal cycling. Three separate plates were run each day to assess annealing temperatures of 55, 57, and 60 C for each assay. The super mixes were formulated for a single dPCR assay per day; for example, on day one of the experiments, S1, S2, and S3 were prepared using the Cow M2 primers and probe. The super mixes required for a single day of experiments were prepared as detailed below. All reagents were thawed from frozen storage (−20 C) and kept on ice during super mix preparation.
3. *A. Super Mix 1 (S1) – 800 nM Forward primer (F), 800 nM Reverse primer (R), 400 nM Probe (P), 24 reactions plus overage.* The following were combined in a 1.7 mL microcentrifuge tube, vortexed for 30 seconds, briefly spun down in a microcentrifuge, and then stored on ice or at 4 C until use: 449 µL molecular-grade water, 250 µL 4X Probe PCR Master Mix from the QIAcuity Probe PCR Kit, 80 µL of 10 µM forward primer solution, 80 µL of 10 µM reverse primer solution, and 16 µL of 25 µM probe solution for a total volume of 875 µL of S1.
4. *B. Super Mix 2 (S2) – 800 nM F, 800 nM R, 250 nM P, 24 reactions plus overage.* The following were combined in a 1.7 mL microcentrifuge tube, vortexed for 30 seconds, briefly spun down in a microcentrifuge, and then stored on ice or at 4 C until use: 455 µL molecular-grade water, 250 µL 4X Probe PCR Master Mix from the QIAcuity Probe PCR Kit, 80 µL of 10 µM forward primer solution, 80 µL of 10 µM reverse primer solution, 10 µL of 25 µM probe solution for a total volume of 875 µL of S2.
5. *C. Super Mix 3 (S3) preparation – 400 nM F, 400 nM R, 250 nM P for 24 reactions plus overage.* The following were combined in a 1.7 mL microcentrifuge tube, vortexed for 30 seconds, briefly spun down in a microcentrifuge, and then stored on ice or at 4 C until use: 535 µL molecular-grade water, 250 µL 4X Probe PCR Master Mix from QIAcuity Probe PCR Kit, 40 µL of 10 µM forward primer solution, 40 µL of 10 µM reverse primer solution, 10 µL of 25 µM probe solution for a total volume of 875 µL of S3.
6. **dPCR reaction setup.** As previously mentioned, during the single-plex assay optimization, three QIAcuity Nanoplate 26k 24-well plates were run each day. Each column of the plate was loaded with a different super mix per the following layout: column 1 - S1 (8 wells), column 2 - S2 (8 wells), column 3 - S3 (8 wells). The dPCR reactions were prepared by pipetting 35 µL of the appropriate super mix into each of eight 0.2 mL tubes in a single PCR strip, followed by adding 5 µL of the 10,000 GC/µL SRM 2917 working solution prepared in Step 1. Three individual strips of 8 PCR tubes, each one containing the appropriate super mix (S1, S2, or S3), were prepared for each experimental run (i.e., single Nanoplate). After the addition of the SRM 2917 working solution, the strip tubes were capped, vortexed for 10 seconds, and then briefly spun down in a microcentrifuge.
7. **Nanoplate 26k 24-well plate loading and sealing.** The entire 40 µL volume from each 0.2 mL PCR tube was transferred into the appropriate well on the Nanoplate, being careful to avoid the introduction of air bubbles (i.e., first-stop pipetting with the pipette tip in contact with the well walls). Once all 24 wells were loaded with reaction mix, the plate was sealed using a Nanoplate Tray, Nanoplate Seal, and the roller provided with the QIAcuity system.
8. **QIAcuity dPCR Experiment Run 1 (55 C annealing temperature).** Once sealed, the Nanoplate was carefully transported into the molecular lab, where the experimental run parameters had already been entered into the QIAcuity Suite Software. For the first experiment, an annealing temperature of 55 C was used with the thermal cycling input as follows: PCR initial heat activation at 95 C for 2 minutes followed by 40 2-step cycles of denaturation at 95 C for 15 seconds and annealing/extension at 55 C for 30 seconds. The Nanoplate barcode was scanned to associate the plate with the correct experimental run in the Suite Software and then carefully loaded into the instrument tray. The run was initiated in the Suite Software and executed per the default settings for partitioning and imaging. Each single-plex dPCR run required approximately 2 hours to complete on the QIAcuity One system. Upon run completion, the dPCR results were exported from the instrument as a CSV file for further analysis.
9. **QIAcuity dPCR Experiment Run 2 (57 C annealing temperature).** For the second single-plex experiment of the day, another Nanoplate 26k 24-well plate was prepared and sealed as described in Steps 3 and 4 above. For the second experiment, an annealing temperature of 57 C was used with the thermal cycling input as follows: PCR initial heat activation at 95 C for 2 minutes followed by 40 2-step cycles of denaturation at 95 C for 15 seconds and annealing/extension at 57 C for 30 seconds. All other aspects of the run were performed as described in Step 5.
10. **QIAcuity dPCR Experiment Run 3 (60 C annealing temperature).** For the third single-plex experiment of the day, another Nanoplate 26k 24-well plate was prepared and sealed as described in Steps 3 and 4. The plate was handled and loaded into the QIAcuity One as previously described, except in this experiment, an annealing temperature of 60 C was used with the thermal cycling input as follows: PCR initial heat activation at 95 C for 2 minutes followed by 40 2-step cycles of denaturation at 95 C for 15 seconds and annealing/extension at 60 C for 30 seconds. Upon run completion, the digital PCR results were exported from the instrument as a CSV file for analysis.

Steps 2 through 7 were repeated daily over six experimental days, running three 26k 24-well Nanoplates for a single dPCR assay each day for a total of 18 plates. These experiments could be planned and executed in a variety of ways, but as an example, our experimental schedule is outlined below:

9 April - Prepare SRM 2917 working solution.

11 April - Cow M2: Nanoplate 1: S1, S2, S3 at 55 C annealing; Nanoplate 2: S1, S2, S3 at 57 C annealing; Nanoplate 3: S1, S2, S3 at 60 C annealing.

12 April - Cow M3 as above.

15 April - Rum2Bac

16 April - Pig2Bac

17 April - Entero1a

25 April - HF183/BacR287

As mentioned previously, the goal of the single-plex experiments was to determine if the separation between positive and negative dPCR partitions was reliable with various primer/probe concentrations and annealing temperatures. Throughout the experiments, we observed reliable and consistent separation between the positive and negative partition clusters for each single-plex assay. In Figure 1, we have provided examples of the QIAcuity 1D scatterplot output for HF183/BacR287 (1A), Pig2Bac (1B), and Entero1a (1C) for nine unique primer/probe concentration and annealing temperature combinations. We observed similar results for the Cow M2, Rum2Bac, and Cow M3 assays (not shown). The separation between negative partitions (grey cluster) and positive partitions (blue cluster) remained reliable for each singleplex assay across all combinations with equivalent thresholding values (red line), yielding comparable results.

**Figure 1.**
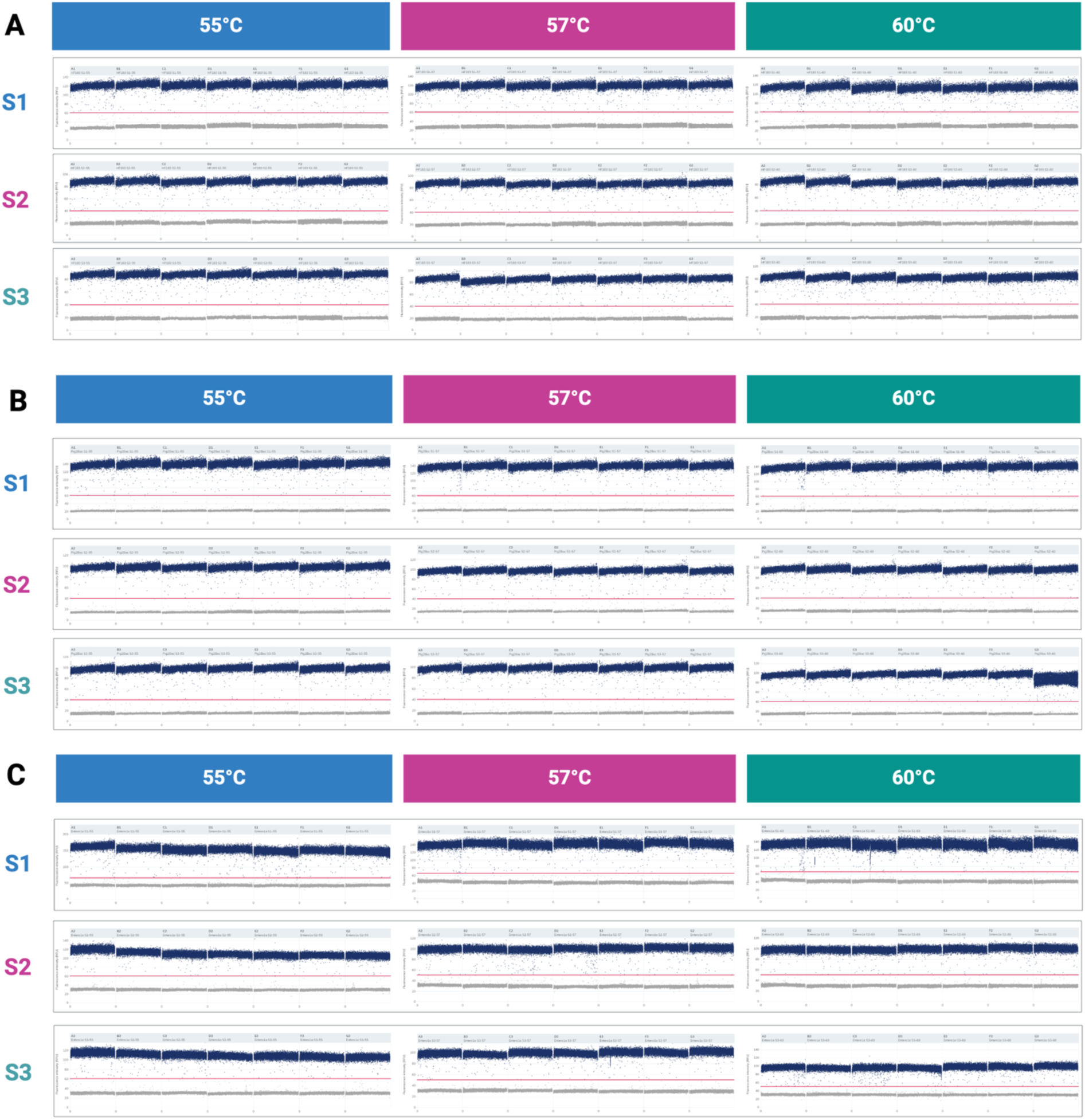
QIAcuity 1D Scatterplots for HF183/BacR287 (A), Pig2Bac (B), and Entero1a (C) assays at nine combinations of annealing temperature (55, 57, 60 C) and primer/probe concentrations (S1, S2, S3).

Based on the results for the single-plex optimization experiments, we proceeded with super mix formulation S3 (400 nM F, 400 nM R, and 250 nM P) to minimize the likelihood of cross talk between channels while returning reliable results during duplex and triplex experiments. Since we observed no difference in cluster separation between annealing temperatures, we also proceeded with an annealing temperature of 60 C (Qiagen’s recommendation) for experiments during duplex and triplex development.

### 3.2 Duplex Assay Optimization

The goal of the duplex optimization experiments was to begin combining MST assays from the anticipated triplex reactions and ensure reliable separation between positive and negative partitions and minimal cross talk between dye channels in duplex format. For this purpose, we created Duplex 1 (D-1) consisting of Cow M2 (green channel) and Cow M3 (crimson channel) and Duplex 2 (D-2) consisting of Pig2Bac (yellow channel) and Entero1a (crimson channel). The two remaining assays, HF183/BacR287 (green channel) and Rum2Bac (orange channel) were combined into Duplex 3 (D-3) even though in the final triplex reactions, they would be separated. The control material for these experiments was the SRM 2917 working solution (∼10,000 GC/µL) created during the single-plex assay optimization (Section 3.1 Step 1). As previously mentioned, based on the observations from the single-plex experiments, all dPCR experiments were run with an annealing temperature of 60 C and with each assay’s reagents added to the reaction per formulation S3.

1. **dPCR duplex super mix preparation.** Duplex super mixes were prepared per formulation S3 (400 nM F, 400 nM R, 250 nM P) for both assays in the duplex. Enough super mix was prepared for 8 reactions plus overage. The following were combined in a 1.7 mL microcentrifuge tube, vortexed for 30 seconds, briefly spun down in a microcentrifuge, and then immediately dispensed as described in Step 2 below: 160.2 µL molecular-grade water, 90 µL 4X Probe PCR Master Mix from the QIAcuity Probe PCR Kit, 14.4 µL of 10 µM assay 1 forward primer solution, 14.4 µL of 10 µM assay 1 reverse primer solution, 3.6 µL of 25 µM assay 1 probe solution, 14.4 µL of 10 µM assay 2 forward primer solution, 14.4 µL of 10 µM assay 2 reverse primer solution, 3.6 µL of 25 µM assay 2 probe working for a total duplex super mix volume of 315 µL.
2. **dPCR reaction setup.** During the duplex assay optimization, a single QIAcuity Nanoplate 26k 8-well plate was run each day – D-1 plate on 29 April, D-2 plate on 30 April, and D-3 plate on 1 May. On each 8-well plate, seven duplex reactions with SRM 2917 control material and one no-template control (NTC) duplex reaction were run. The dPCR reactions were prepared by pipetting 35 µL of the day’s super mix into each of eight 0.2 mL tubes in a single PCR strip, followed by adding 5 µL of the 10,000 GC/µL SRM 2917 working solution into wells A to G and 5 µL of molecular-grade water into well H (NTC). After the addition of the SRM 2917 working solution (or molecular-grade water for NTCs), the strip tubes were capped, vortexed for 10 seconds, and then briefly spun down in a microcentrifuge.
3. **Nanoplate 26k 8-well plate loading and sealing.** The entire 40 µL volume from each 0.2 mL PCR tube was transferred into the appropriate well on the Nanoplate, being careful to avoid the introduction of air bubbles. Once all 8 wells were loaded with the appropriate reaction mix, the plate was sealed using a Nanoplate Tray, Nanoplate Seal, and the roller provided with the QIAcuity system.
4. **QIAcuity dPCR Experiment Run.** Once sealed, the 8-well Nanoplate was carefully transported into the molecular lab, where the experimental run parameters had already been entered into the QIAcuity Suite Software. An annealing temperature of 60 C was used with the thermal cycling input as follows: PCR initial heat activation at 95 C for 2 minutes followed by 40 2-step cycles of denaturation at 95 C for 15 seconds and annealing/extension at 60 C for 30 seconds. To avoid oversaturation during plate imaging, the gain and exposure were adjusted from the default values to achieve satisfactory results (Note 5). Otherwise, each duplex dPCR experiment run was performed as previously described in Section 3.1 Step 7.

Just as the single-plex experiments, the overall goal of the duplex experiments was to determine if the separation between positive and negative dPCR partitions was reliable for both assays in each duplex while cross talk between channels was minimal. In Figure 2, we have provided examples of the QIAcuity 1D scatterplot output for single-plex (left column) versus duplex (right column) format for each MST assay. The separation between negative partitions (grey cluster) and positive partitions (blue cluster) remained consistent for each duplex pair, with thresholding (red line) at similar levels. Throughout the duplex experiments, we observed reliable and consistent separation between the positive and negative partition clusters for each assay in duplex format compared to single-plex format. The number of partitions falling in the intermediate fluorescence intensity range (referred to as “rain”) was comparable between single-plex and duplex formats, suggesting cross talk did not affect the results.

**Figure 2.**
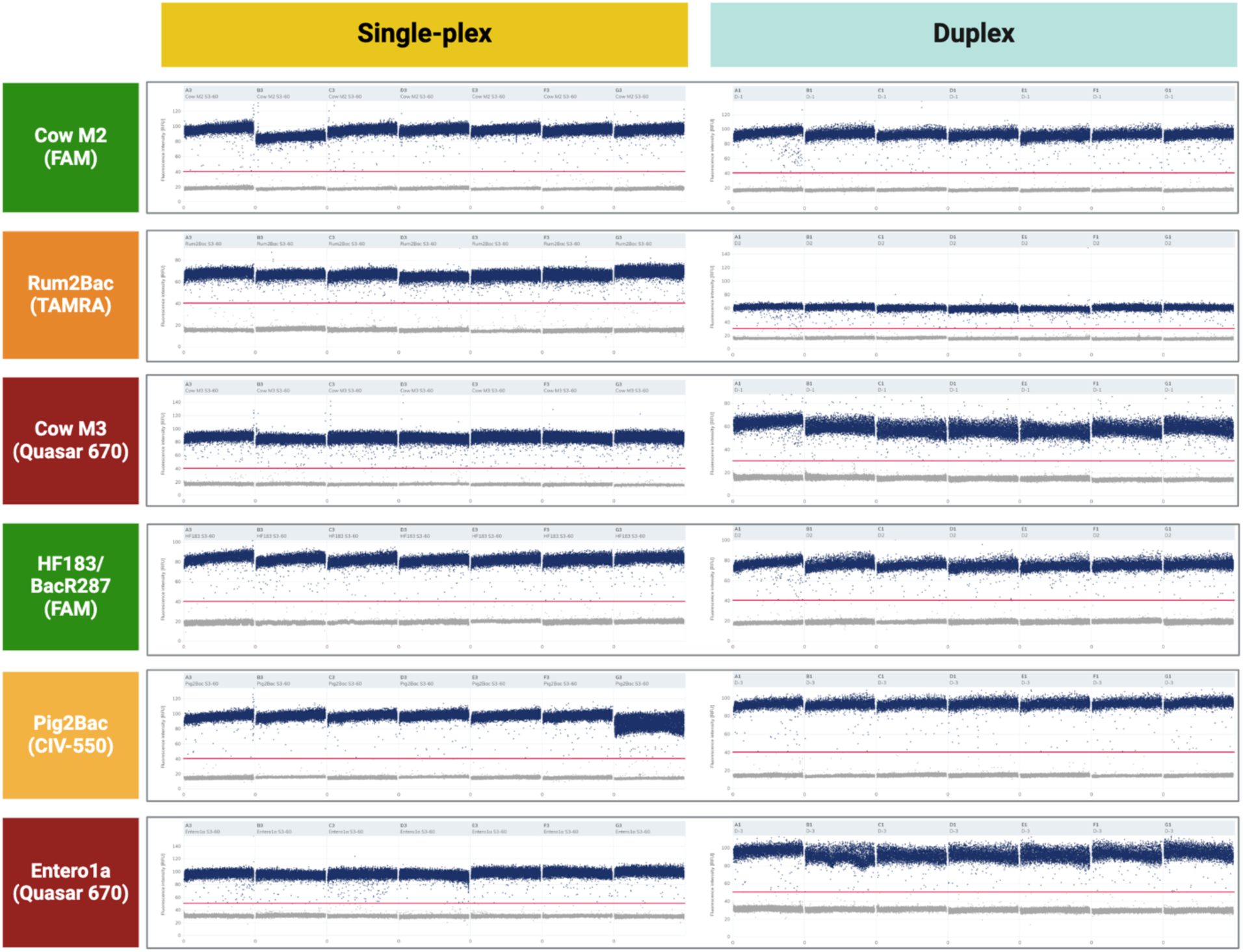
QIAcuity 1D scatterplots for each dPCR MST assay in single-plex (left column) versus duplex (right column) format.

Quantitative results from the duplex experiments are shown in Figure 3. Owing to the long working life of the SRM 2917 working solution (22 days), we observed some decay from the initial 10,000 GC/µL concentration. Within each duplex, individual MST assays yielded statistically similar concentrations (*p* > 0.05 Wilcoxon signed rank tests). However, between experimental runs (i.e., D-1, D-2, D-3), the estimated SRM 2917 varied from a mean of 3.52 log_10_ GC/µL by the D-1 assays to a mean of 3.41 log_10_ GC/µL by the D-2 assays. Nonetheless, the estimated SRM 2917 concentrations across all three duplexes were within 0.11 log_10_ GC/µL of one another.

**Figure 3.**
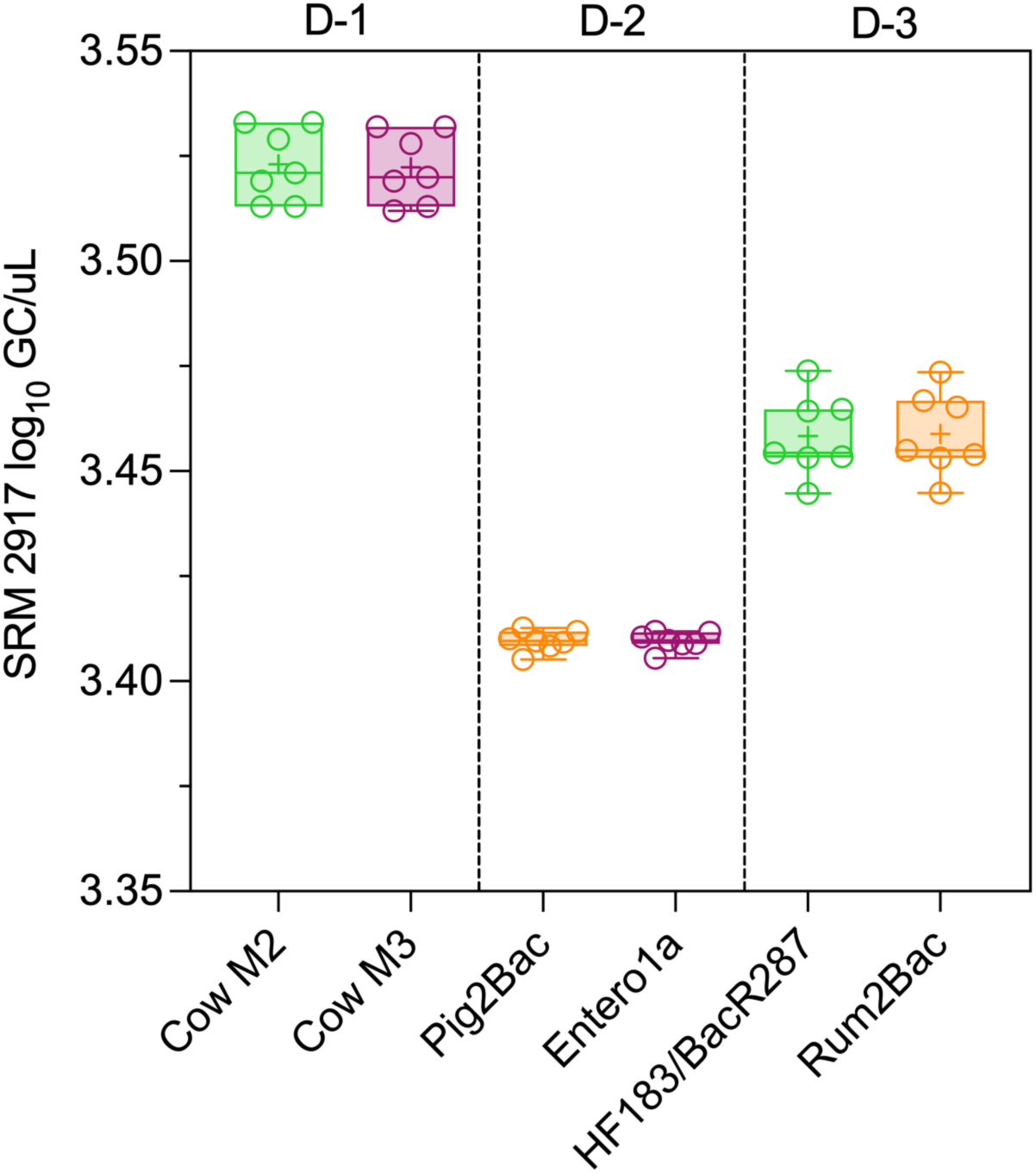
SRM 2917 concentration as estimated by individual MST assays performed in three duplex format reactions (D-1, D-2, D-3) with seven technical replicates per duplex.

### 3.3 Triplex Assay Optimization Fixed Concentration

Based on the successful performance of the MST assays in duplex format, we proceeded to the final triplex format by adding Rum2Bac to D-1 to create Triplex 1 (Cow M2, Rum2Bac, Cow M3) and adding HF183/BacR287 to D-2 to create Triplex 2 (HF183/Rum2Bac, Pig2Bac, Entero1a). For the dPCR triplex reaction super mixes, we continued to use formulation S3 for each individual assay, and we maintained an annealing temperature of 60 C. To assess the performance of the triplex dPCR reactions, we created a fresh batch of SRM 2917 solution at a fixed concentration.

1. **SRM 2917 working solution preparation.** Based on the anticipated sample size for the triplex experiments, a 300 µL SRM 2917 working solution at a final concentration of approximately 10,000 GC/µL was prepared by pipetting 294.5 µL of molecular-grade water into a 1.5 mL LoBind centrifuge tube followed by 5.5 µL of SRM 2917 Level 6 stock solution (certified by NIST at 541,286 +/- 7,700 GC/µL). The SRM 2917 working solution was stored at 4 C when not in use across the two experiments in 25 calendar days. Prior to any subsampling for dPCR experiments, the tube was vortexed for 30 seconds and briefly spun down in a microcentrifuge.
2. **dPCR triplex super mix preparation.** Triplex super mixes were prepared per formulation S3 (400 nM F, 400 nM R, 250 nM P) for each assay in the triplex. Enough triplex super mix was prepared for six reactions plus overage. The following were combined in a 1.7 mL microcentrifuge tube, vortexed for 30 seconds, briefly spun down in a microcentrifuge, and then immediately dispensed as described in Step 3 below: 99.4 µL molecular-grade water, 70 µL 4X Probe PCR Master Mix from the QIAcuity Probe PCR Kit, 11.2 µL of 10 µM assay 1 forward primer solution, 11.2 µL of 10 µM assay 1 reverse primer solution, 2.8 µL of 25 µM assay 1 probe solution, 11.2 µL of 10 µM assay 2 forward primer solution, 11.2 µL of 10 µM assay 2 reverse primer solution, 2.8 µL of 25 µM assay 2 probe solution and 11.2 µL of 10 µM assay 3 forward primer solution, 11.2 µL of 10 µM assay 3 reverse primer solution, 2.8 µL of 25 µM assay 3 probe solution for a total triplex super mix volume of 245 µL. Single-plex reaction mixes for each MST assay were prepared per formulation S3 for six reactions (149.8 µL molecular-grade water, 70 µL 4X Probe PCR Master Mix from the QIAcuity Probe PCR Kit, 11.2 µL of 10 µM forward primer solution, 11.2 µL of 10 µM reverse primer solution, 2.8 µL of 25 µM probe solution for a total single-plex super mix volume of 245 µL). For each assay, enough single-plex super mix was prepared for six reactions plus overage.
3. **dPCR reaction setup.** During the triplex SRM 2917 fixed concentration experiments, a single QIAcuity Nanoplate 26k 24-well plate was run each day – Triplex 1 on 8 July and Triplex 2 on 2 August. On each 24-well plate, five triplex technical replicates with SRM 2917 (*n* = 5), five single-plex technical replicates with SRM 2917 per individual assay (*n* = 15), and one NTC per triplex and single-plex reaction (*n* = 4) for a total of 24 wells. Running both triplex and single-plex reactions on a single plate allows for the most reliable comparison of performance between the two formats. The dPCR reactions were prepared by pipetting 35 µL of the appropriate super mix into each of six 0.2 mL tubes in a single PCR strip, followed by adding 5 µL of the 10,000 GC/µL SRM 2917 working solution five tubes and 5 µL of molecular-grade water into a single tube for the five technical replicates and one NTC for each assay. After the addition of the SRM 2917 working solution (or molecular-grade water for NTCs), the strip tubes were capped, vortexed for 10 seconds, and then briefly spun down in a microcentrifuge.
4. **Nanoplate 26k 24-well plate loading and sealing.** The entire 40 µL volume from each 0.2 mL PCR tube was transferred into the appropriate well on the Nanoplate, being careful to avoid the introduction of air bubbles. Once all 24 wells were loaded with the appropriate reaction mix, the plate was sealed using a Nanoplate Tray, Nanoplate Seal, and the roller provided with the QIAcuity system.
5. **QIAcuity dPCR Experiment Run.** Once sealed, the 24-well Nanoplate was carefully transported into the molecular lab, where the experimental run parameters had already been entered into the QIAcuity Suite Software. An annealing temperature of 60 C was used with the thermal cycling input as follows: PCR initial heat activation 95 C for 2 minutes followed by 40 2-step cycles of denaturation 95 C for 15 seconds and annealing/extension 60 C for 30 seconds. The dPCR experimental run for each triplex plate was performed as per Section 3.2 Step 4.

Across both the Triplex 1 and Triplex 2 experiments (Figure 4), the performance of the triplex (T) format versus single-plex (S) format for quantifying SRM 2917 was statistically comparable (*p* > 0.05). The range of SRM 2917 concentrations estimated for the five technical replicates of each MST assay was within 0.03 log_10_ of the expected value (4.00 log_10_ GC/µL) for Triplex 1 (left panel) and 0.02 log_10_ for Triplex 2 (right panel).

**Figure 4.**
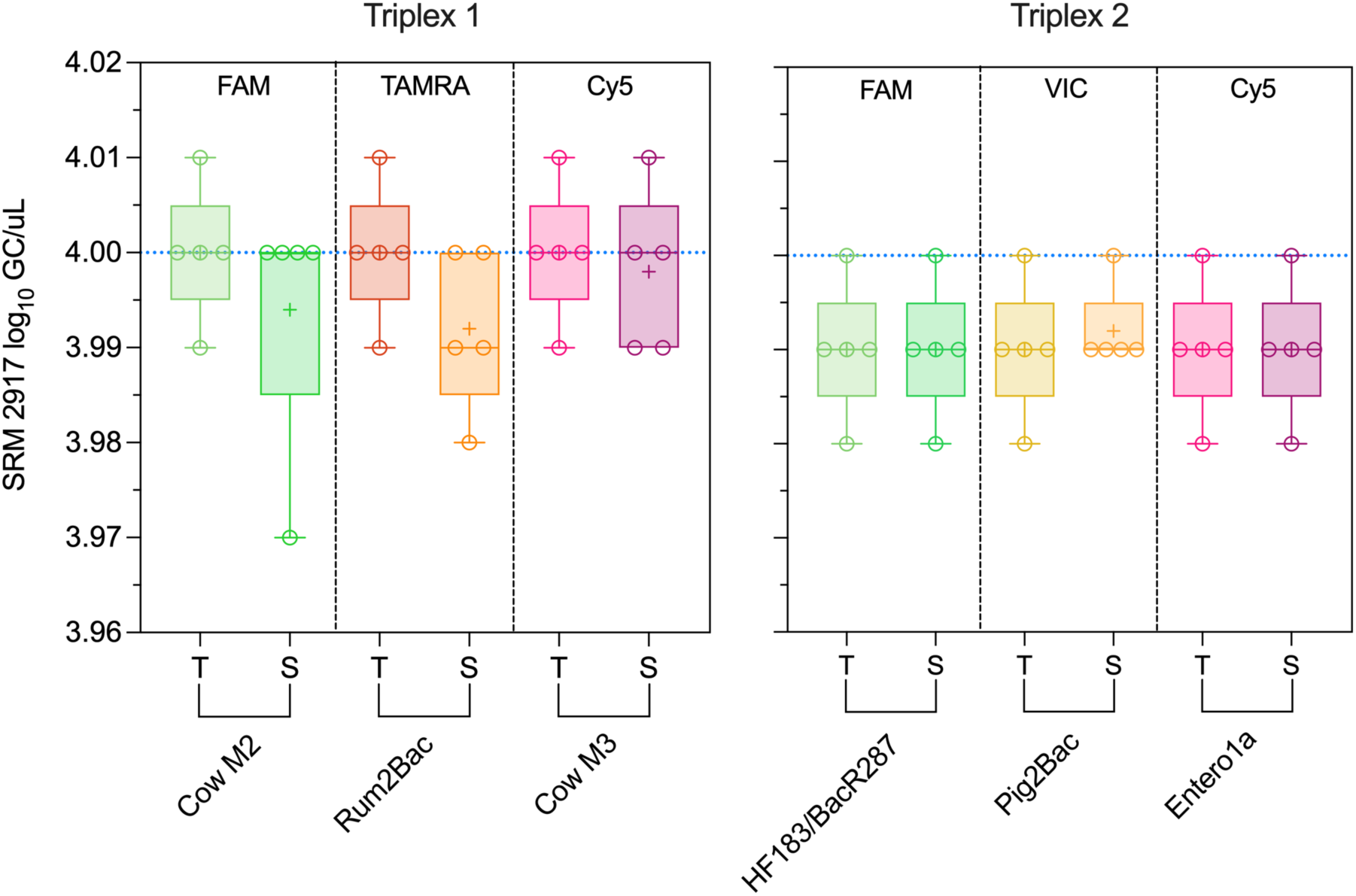
Quantification of the SRM 2917 control material by two separate triplex reactions (Triplex 1, Triplex 2) yielded statistically similar (*p* > 0.05) quantities of SRM 2917 by each assay in triplex format (T) compared to the same assay in single-plex (S) format.

### 3.4 Triplex Assay Optimization Concentration Gradient

To assess the performance of the triplex assays further, we also completed triplex versus single-plex comparison experiments using a concentration gradient of SRM 2917. Using 26k Nanoplate 24-well plates, we performed experiments in a similar format to Section 3.3, except during this experimental round, each assay in each format (T, S) was tested in two technical replicates against three dilution levels for SRM 2917. For dPCR triplex and single-plex reaction super mixes, we continued to use formulation S3, and we maintained an annealing temperature of 60 C.

1. **SRM 2917 gradient working solution preparation.** Based on the anticipated sample size for the triplex gradient experiments, a concentration gradient of SRM 2917 working solutions was prepared. The first-level dilution of 200 µL at 250 GC/µL was made by mixing 9.40 µL SRM 2917 Level 4 (NIST certified value 5,314 GC/µL +/- 71 GC/µL) and 190.60 µL molecular grade water in a 1.5 mL LoBind centrifuge tube. The second-level dilution of 250 µL at 50 GC/µL was prepared by mixing 50 uL of the first-level working solution and 200 µL of molecular-grade water in a 1.5 mL LoBind centrifuge tube. The third-level dilution of 150 µL at 25 GC/µL was prepared by mixing 75 µL of the second-level working solution and 75 µL of molecular-grade water in a LoBind centrifuge tube. The SRM 2917 concentration gradient working solutions were stored at 4 C when not in use across the two experiments in 30 calendar days. Prior to any subsampling for dPCR experiments, the tube was vortexed for 30 seconds and briefly spun down in a centrifuge.
2. **dPCR triplex super mix preparation.** Triplex super mixes were prepared per formulation S3 (400 nM F, 400 nM R, 250 nM P) for each assay in the triplex. Enough triplex super mix was prepared for six reactions plus overage as described in Section 3.3 Step 2. Single-plex reaction mixes for each MST assay were also prepared per formulation S3 as detailed in Section 3.3 Step 2. For each assay, enough single-plex super mix was prepared for six reactions plus overage.
3. **dPCR reaction setup.** During the triplex SRM 2917 concentration gradient experiments, a single QIAcuity Nanoplate 26k 24-well plate was run each day – Triplex 1 on 9 July and Triplex 2 on 8 August. Each 24-well plate included two triplex technical replicates for each SRM 2917 dilution level (*n* = 6) and two single-plex technical replicates for each SRM 2917 dilution level per individual assay (*n* = 18, 6 per assay by 3 assays) for a total of 24 wells. The dPCR reactions were prepared by pipetting 35 µL of the appropriate super mix into each of six 0.2 mL tubes in a single PCR strip, followed by adding 5 µL of the appropriate SRM 2917 dilution level. After the addition of the SRM 2917 working solution, the PCR strip tubes were capped, vortexed for 10 seconds, and then briefly spun down in a microcentrifuge.
4. **Nanoplate 26k 24-well plate loading and sealing.** The entire 40 µL volume from each 0.2 mL PCR tube was transferred into the appropriate well on the Nanoplate, and the plate was sealed as previously described.
5. **QIAcuity dPCR Experiment Run.** Once sealed, the 24-well Nanoplate was carefully transported into the molecular lab, where the experimental run parameters had already been entered into the QIAcuity Suite Software. An annealing temperature of 60 C was used with the thermal cycling input as follows: PCR initial heat activation 95 C for 2 minutes followed by 40 2-step cycles of denaturation 95 C for 15 seconds and annealing/extension 60 C for 30 seconds. The dPCR experimental run for each triplex plate was again performed as per Section 3.2 Step 4.

As shown in Figure 5, quantification of the SRM 2917 control material along a concentration gradient (2.40, 1.70, 1.40 log_10_ GC/µL) by Triplex 1 (panel A) and Triplex 2 (panel B) yielded similar quantities of SRM 2917 compared to the same assay in single-plex (S) (maximum difference between T/S pairs < 0.15 log_10_ GC/µL). The maximum difference between the mean concentration estimated from two technical replicates for each triplex assay and the expected value across the entire concentration gradient was 0.13 log_10_ GC/µL. When considering the QIAcuity 1D scatterplots, Triplex 1 (Figure 6) and Triplex 2 (Figure 7) provided reasonable separation between the positive (blue) and negative (grey) partitions such that equivalent thresholding (red line) for triplex (left column) and single-plex (right column) reactions yielded similar quantitative results.

**Figure 5.**
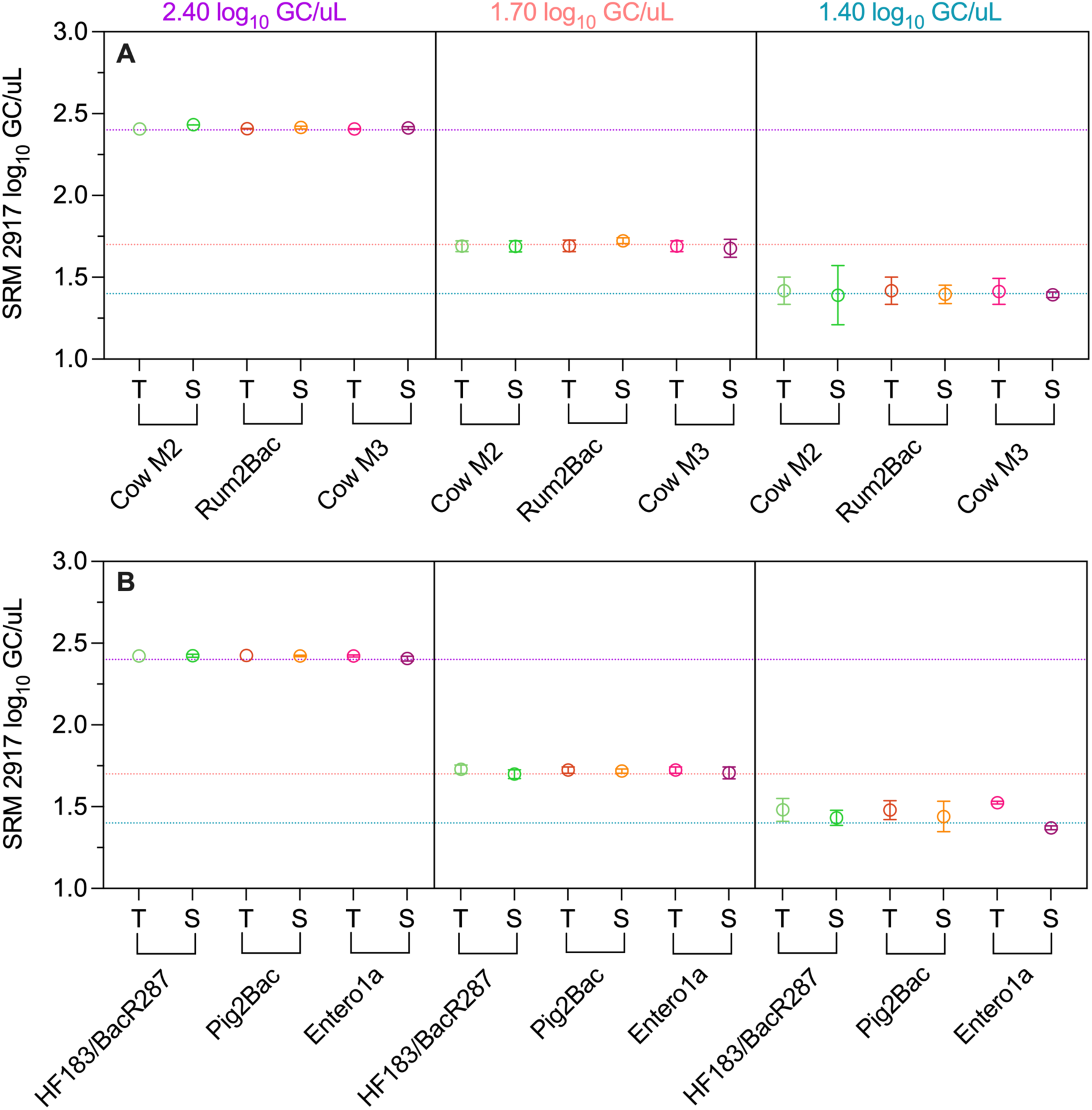
Quantification of the SRM 2917 control material along a concentration gradient (2.40, 1.70, 1.40 log_10_ GC/µL) by Triplex (T) 1 (panel A) and Triplex (T) 2 (panel B) compared to single-plex (S) format results for each MST assay.

**Figure 6.**
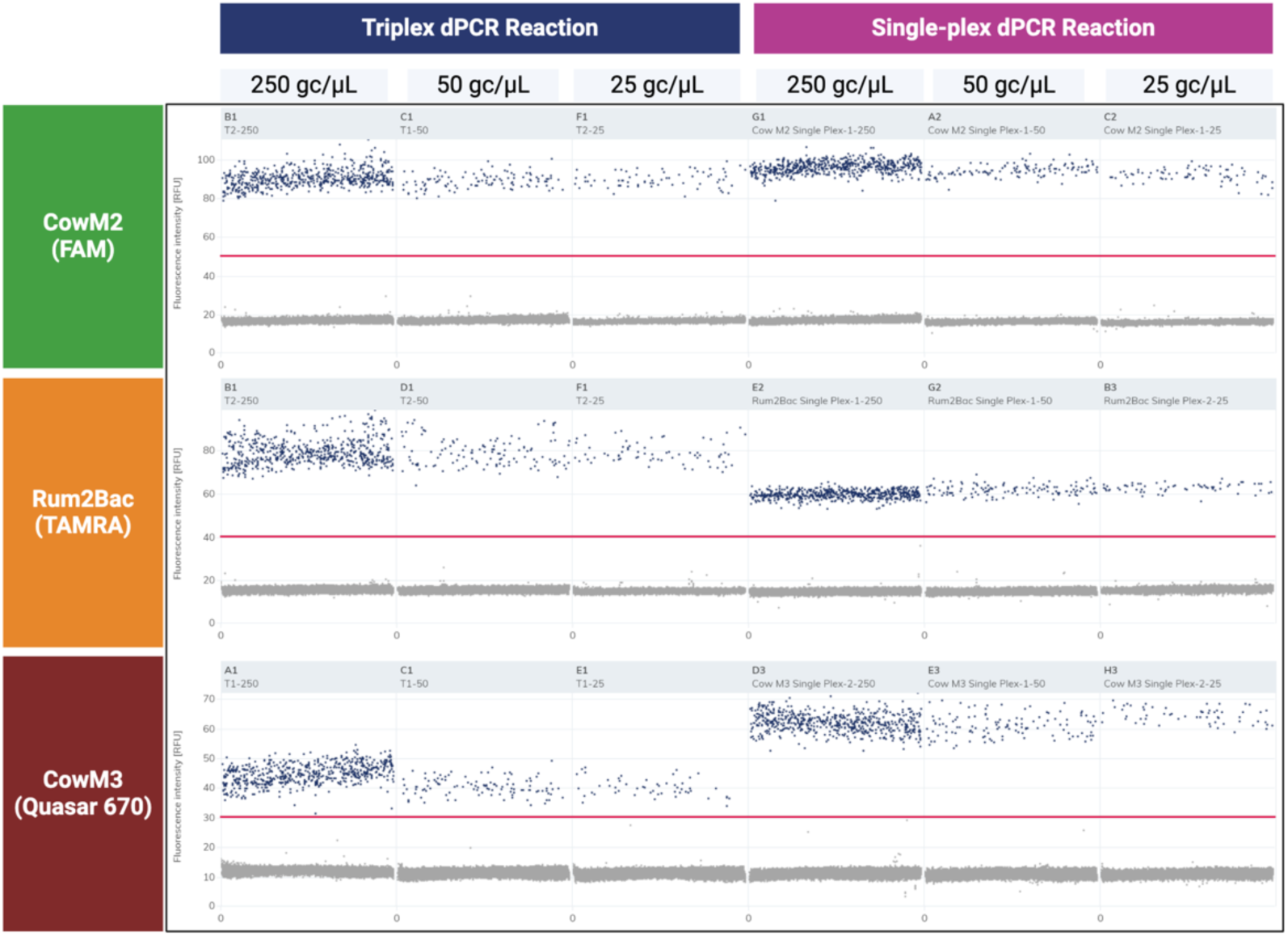
QIAcuity 1D scatterplots for triplex versus single-plex format dPCR reactions for Triplex 1 (CowM2, Rum2Bac, CowM3) along a concentration gradient of SRM 2917 control material (250 GC/µL, 50 GC/µL, 25 GC/µL) as viewed on the QIAcuity software.

**Figure 7.**
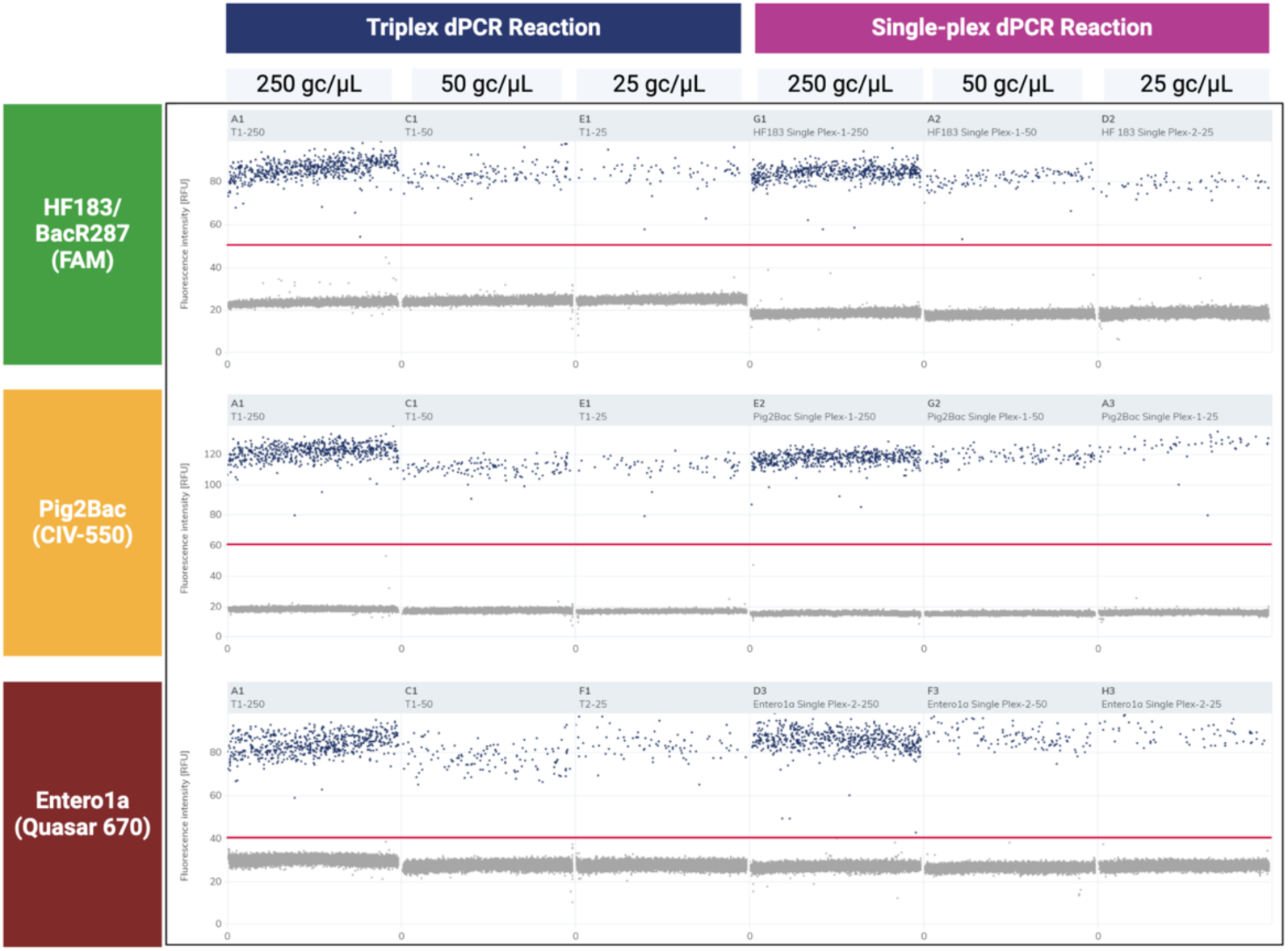
QIAcuity 1D scatterplots for triplex versus single-plex dPCR reactions for Triplex 2 (HF183/BacR287, Pig2Bac, Entero1a) along a concentration gradient of SRM 2917 control material (250 GC/µL, 50 GC/µL, 25 GC/µL) as viewed on the QIAcuity software.

### 3.5 Triplex Testing of Archived Surface Water Extracts

Having established the performance of the triplex assays for quantifying an ideal standard reference material, we wished to challenge them against purified nucleic acids derived from actual surface water samples. The six nucleic acid samples were produced during a previous water quality study (Section 2.3) and had been tested in single-plex format by QIAcuity using some of the MST assays before being stored at -80 C. We also included two additional surface water samples that had not been previously tested. Triplex reaction mixes were prepared for 8 reactions plus overage per formulation S3 (127.8 µL molecular-grade water, 90 µL 4X Probe PCR Master Mix from the QIAcuity Probe PCR Kit, 14.4 µL of 10 µM assay 1 forward primer solution, 14.4 µL of 10 µM assay 1 reverse primer solution, 3.6 µL of 25 µM assay 1 probe solution, 14.4 µL of 10 µM assay 2 forward primer solution, 14.4 µL of 10 µM assay 2 reverse primer solution, 3.6 µL of 25 µM assay 2 probe solution and 14.4 µL of 10 µM assay 3 forward primer solution, 14.4 µL of 10 µM assay 3 reverse primer solution, 3.6 µL of 25 µM assay 3 probe solution for a total triplex super mix volume of 315 µL). A single QIAcuity Nanoplate 26k 8-well plate was run for each triplex, consisting of two triplex technical replicates for each of the four water samples. Triplex 1 was tested against two samples that had previously tested positive for Rum2Bac and Cow M3 in single-plex format (EA, WA) and two unknown samples (NR4, YW4). Triplex 2 was tested against two samples that had previously tested positive for HF183/BacR287 and Entero1a by single-plex assays (F1021, COM3), and the same two unknown samples as used for Triplex 1 (NR4, YW4). The dPCR reactions were prepared by pipetting 35 µL of the appropriate super mix into each of eight 0.2 mL tubes in a single PCR strip, followed by adding 5 µL of the purified DNA from the appropriate sample. After the addition of the sample DNA, the strip tubes were capped, vortexed for 10 seconds, and then briefly spun down in a microcentrifuge. The entire 40 µL volume from each 0.2 mL PCR tube was transferred into the Nanoplate and the plate was sealed per the manufacturer’s protocol as previously described. An annealing temperature of 60 C was used and the dPCR experimental run for each triplex plate was performed as described in Section 3.2 Step 4.

As shown in Figure 8 for Triplex 1, samples that had previously tested positive (EA, WA) for Rum2Bac and CowM3 in single-plex (S) format also tested positive for each MST target in the triplex (T) format. The concentrations estimated by Triplex 1 were comparable to those estimated in single-plex for both Rum2Bac (∼4.5 log_10_ GC/100 mL) and Cow M3 (∼4 log_10_ GC/100 mL). The water samples, which were seeded with cow manure, also tested positive for Cow M2 in one of two technical replicates for each sample, although previous single-plex results were not available for Cow M2. The two unknown water samples (NR4 and YW4) were non-detect for all three MST targets, which are associated with cows and ruminants. While there are no single-plex results available for comparison, the suspected source of fecal contamination in these two waterbodies is human sewage from failing on-site sewer systems, so negativity for ruminant MST markers is plausible.

**Figure 8.**
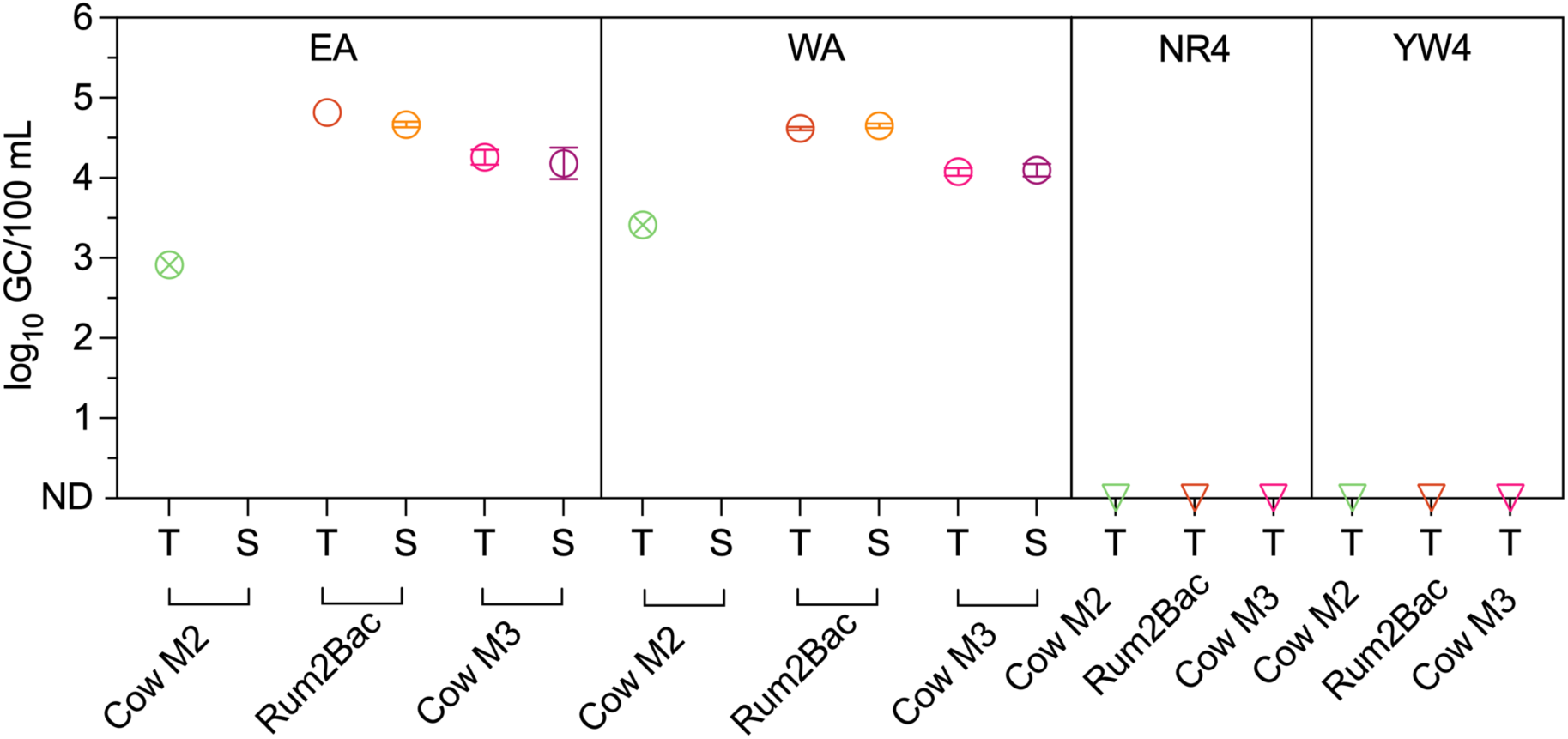
MST dPCR Triplex 1 (T) results from dPCR duplicates for two archived samples (EA, WA) compared to previously generated single-plex (S) results for Rum2Bac and Cow M3. MST dPCR triplex results from dPCR duplicates for two archived samples (NR4, YW4) not previously tested in single-plex format. Inverted triangles indicate non-detection in both replicates; circles with x’s denote one detection among two technical replicates at the concentration estimated from that single detection; open circles with error bars denote the mean and standard deviation for samples testing positive in both replicates; if open circles are shown without error bars then the standard deviation is too small to be displayed.

For Triplex 2, samples that had previously tested positive for HF183/BacR287 and Entero1a by single-plex (S) format also tested positive in triplex format (T), as indicated in Figure 9. For sample F1021, the concentration of HF183/BacR287 by Triplex 2 was about 0.6 log_10_ GC/100 mL higher than by single-plex format. While for COM3, HF183/BacR287 was only detected in a single technical replicate in the triplex format, while both single-plex technical replicates were previously positive. However, the estimated concentration (∼1.5 to 2.0 log_10_ GC/100 mL) was comparable. For Entero1a all technical replicates in both triplex and single-plex format were positive with similar concentration estimates (3.5 to 4.0 log_10_ GC/100 mL) in both formats. Pig2Bac, an MST marker for pigs, was detected in a single technical replicate for COM3, but single-plex results are not available for comparison. For the unknown samples (NR4, YW4), Entero1a was detected by triplex format in all technical replicates with an estimated concentration of 3.0 to 3.5 log_10_ GC/100 mL. Although single-plex results from these specific samples are not available, this abundance is consistent with other Entero1a testing of samples from these waterbodies (data not shown). Pig2Bac was not detected in either sample. HF183/BacR287 was detected in all technical replicates from NR4 and YW4 at concentrations of approximately 3 log_10_ GC/100 mL across both samples. These results are consistent with the suspected source of fecal contamination for these waterbodies – human sewage from failing on-site sewer systems.

**Figure 9.**
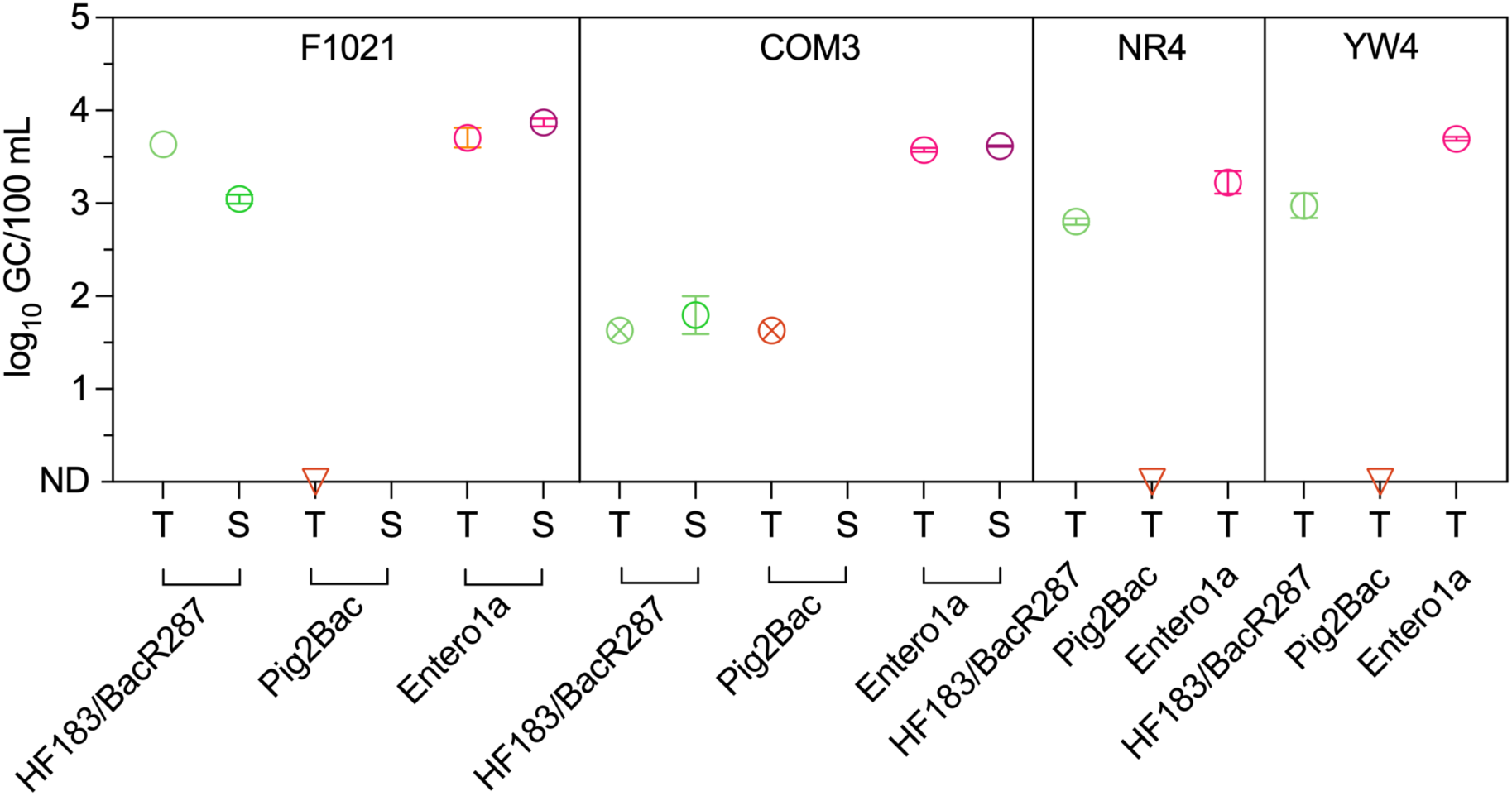
MST dPCR triplex (T) results from dPCR technical replicates for two archived samples (F1021, COM3) compared to previously generated single-plex (S) results for HF183/BacR287 and Entero1a. MST dPCR triplex results from dPCR duplicates for two archived samples (NR4, YW4) not previously tested in single-plex format. Inverted triangles indicate non-detection in both replicates; circles with x’s denote one detection among two technical replicates at the concentration estimated from that single detection; open circles with error bars denote the mean and standard deviation for samples testing positive in both replicates; if open circles are shown without error bars then the standard deviation is too small to be displayed.

### 3.6 Conclusions and Final Recommendations

Here, we have described a detailed and multi-phased protocol to develop and validate two triplex assays for MST in surface waters via dPCR performed on the QIAcuity One system. Although our protocol is specific to the assays, standard reference materials, and sample materials tested, the approach could easily be extended and amended to develop other multiplex assays for dPCR – up to 5-plex on the QIAcuity One system. We conservatively elected to develop our MST assays as two triplexes so that we could skip dye channels on the QIAcuity; however, given our success with the triplex format, we now plan to further test additional assays in the skipped channels for each triplex. Based on our experience, crucial variables in the successful empirical optimization include forward and reverse primer compatibilities (i.e., primer dimers) and concentrations, probe compatibilities and concentration, annealing temperature compatibility, and reliable quantitative standard materials. Each of the triplex MST dPCR reactions developed in the current work quantified a standard reference material, at both fixed concentration and across a concentration gradient down to 25 GC/µL of added template at a statistically comparable measurement to the single-plex MST assays. The triplex assays also provided MST maker concentration estimates that were comparable to previous single-plex MST results for actual surface water samples. It should be noted that each of the molecular assays used in this work were previously optimized and developed for MST purposes and that such considerations have been beyond the scope of our current consideration. While *in silico* analysis is always advisable for new dPCR/qPCR assay development, utimately, in our experience, optimization must be performed in an empirical fashion with our protocols described here as a useful example for others.

## 4 Notes

1. Based on conversations with other QIAcuity users, we opted to skip dye channels to minimize the likelihood of crosstalk during triplex dPCR reactions. We consulted the Biosearch Technologies Flourophore and BHQ selection chart to select probe and quencher combinations based on the five dye channels on the QIAcuity system (***20***). The anticipated spectral overlay for probe selections can be visualized using Biosearch Technologies’ Spectral Overlay Tool (***21***).
2. Biosearch Technologies has very useful resources to consult while developing singleplex and multiplex dPCR assays, including information on: Considerations for multiplexing (***22***), Fluorophore/Quencher combinations (***23***), and Quenching Mechanisms (***24***).
3. A quantitative standard is extremely useful for dPCR assay validation and optimization. In this case, SRM2917 is provided by NIST in a 6-level dilution series from 5 to 500,000 copies per µL (***25***). Upon receipt, we quantified the lowest dilution level (Level 1) via dPCR (QIAcuity) using the HF183/BacR287 assay within one copy per µL of the certified value prior to proceeding with the experiments we describe here. If no quantitative standards are used, we recommend using dPCR rather than mass-based estimates to establish the control stock solution titer. In our experience, mass-based estimates of the copy number frequently exceed the copy number that can be verified by dPCR.
4. As previously mentioned, minimizing cross talk between dye channels is one of the key considerations for developing reliable multiplex dPCR assays. In our experience and the experience of others, the manufacturer-recommended concentrations for primers and probes on the QIAcuity system are higher than necessary and may create the opportunity for cross talk. Therefore, we experimented with primer and probe concentrations that were lower than the manufacturer’s recommendations.
5. Optimize imaging parameters for each dye channel by progressively lowering exposure duration and gain settings. Begin with the brightest channel and decrease these values for subsequent, less intense channels. For example: FAM (brightest): 500 ms exposure, gain of 6; TAMRA (medium): 400 ms exposure, gain of 5; Cy5 (dimmest): 325 ms exposure, gain of 4. This stepwise reduction in settings ensures appropriate signal capture across channels with varying fluorescence intensities.

